# Individual differences in dopamine are associated with reward discounting in clinical groups but not in healthy adults

**DOI:** 10.1101/383810

**Authors:** Jaime J. Castrellon, Kendra L. Seaman, Jennifer L. Crawford, Jacob S. Young, Christopher T. Smith, Linh C. Dang, Ming Hsu, Ronald L. Cowan, David H. Zald, Gregory R. Samanez-Larkin

**Affiliations:** Department of Psychology and Neuroscience, Duke University; Center for Cognitive Neuroscience, Duke University; Center for the Study of Aging and Human Development, Duke University; Department of Psychology, Yale University; Department of Psychology, Vanderbilt University; Haas School of Business, University of California Berkeley; Department of Psychiatry and Behavioral Sciences, Vanderbilt University School of Medicine; Department of Radiology and Radiological Sciences, Vanderbilt University Medical Center

**Keywords:** decision making, delay discounting, probability, effort, dopamine, PET

## Abstract

Some people are more willing to make immediate, risky, or costly reward-focused choices than others, which has been hypothesized to be associated with individual differences in dopamine (DA) function. In two studies using PET imaging, one empirical (Study 1: N=144 males and females across 3 samples) and one meta-analytic (Study 2: N=307 across 12 samples), we sought to characterize associations between individual differences in DA and time, probability, and physical effort discounting in human adults. Study 1 demonstrated that individual differences in DA D2-like receptors were not associated with time or probability discounting of monetary rewards in healthy humans, and associations with physical effort discounting were inconsistent across adults of different ages. Meta-analytic results for temporal discounting corroborated our empirical finding for minimal effect of DA measures on discounting in healthy individuals, but suggested that associations between individual differences in DA and reward discounting depend on clinical features. Addictions were characterized by negative correlations between DA and discounting but other clinical conditions like Parkinson’s Disease, obesity, and ADHD were characterized by positive correlations between DA and discounting. Together the results suggest that trait differences in discounting in healthy adults do not appear to be strongly associated with individual differences in D2-like receptors. The difference in meta-analytic correlation effects between healthy controls and individuals with psychopathology suggests that individual difference findings related to DA and reward discounting in clinical samples may not be reliably generalized to healthy controls, and vice-versa.

**Significance Statement:** Decisions to forgo large rewards for smaller ones due to increasing time delays, uncertainty, or physical effort have been linked to differences in dopamine (DA) function, which is disrupted in some forms of psychopathology. It remains unclear whether alterations in DA function associated with psychopathology also extend to explaining associations between DA function and decision making in healthy individuals. We show that individual differences in dopamine D2 receptor availability are not consistently related to monetary discounting of time, probability, or physical effort in healthy individuals across a broad age range. By contrast, we suggest that psychopathology accounts for observed inconsistencies in the relationship between measures of dopamine function and reward discounting behavior.

## Introduction

Discounting is a natural phenomenon that describes the tendency to devalue rewards that are relatively delayed, uncertain, or require more effort than sooner, more certain, or less effortful ones. Individual differences in discounting in humans have been hypothesized to be strongly related to individual differences in dopamine (DA) function. Studies of human and non-human animals have reported that pharmacological effects on DA D2-like receptors alter discounting (Salamone et al., 1996; St Onge et al., 2010; Koffarnus et al., 2011; Weber et al., 2016). Specifically, D2-like receptors are believed to regulate decisions to inhibit impulsive actions (Frank, 2005; Ghahremani et al., 2012; Robertson et al., 2015) like choosing smaller-sooner/more-likely/less-effortful rewards. However, studies of the transient manipulation of the DA system do not clarify whether more persistent individual differences in decision making are also primarily mediated by differences in DA D2-like receptor expression.

Multiple studies have reported links between discounting behavior and forms of psychopathology that are associated with alteration in striatal DA function including: drug addiction (MacKillop et al., 2011; Amlung et al., 2017), obesity (Amlung et al., 2016), schizophrenia and bipolar disorder (Ahn et al., 2011), attention-deficit/hyperactivity disorder (ADHD) (Amlung et al., 2016), and Parkinson’s disease (PD) (Kaasinen and Vahlberg, 2017). While these studies suggest a common involvement of DA in discounting in disease, it leaves open questions about specific features and clinical range of influence between DA and discounting behavior.

Only a few studies have directly assessed associations between trait-like individual differences in DA function and discounting behavior. Several recent studies using positron emission tomography (PET) suggest that reduced availability of DA receptors contributes to greater discounting (See Table 1 for a summary of dopamine PET studies of reward discounting). However, many existing studies are limited by small sample sizes (Button et al., 2013), a focus on only temporal discounting (Crunelle et al., 2014; Ballard et al., 2015; Cho et al., 2015; Joutsa et al., 2015; Oberlin et al., 2015; Smith et al., 2016) or a mixture of decision features which may or may not be dissociable (Treadway et al., 2012), use of radiotracers with limited visibility outside the striatum (e.g., [11C]raclopride), or assessment of individuals with psychopathology (that vary in DA and other neuromodulatory functions) (Crunelle et al., 2014; Ballard et al., 2015; Eisenstein et al., 2015; Joutsa et al., 2015; Oberlin et al., 2015). Although prior PET studies have largely focused on the striatum, DA neurons in the midbrain also project to the amygdala, hippocampus, thalamus, anterior cingulate, insula, and frontal and parietal lobes (Bjorklund et al., 1978; Berger et al., 1991). Accordingly, there may be subtle differences in how DA function uniquely accounts for different types of discounting across the brain in individuals who vary in DA status.

**Table 1.**
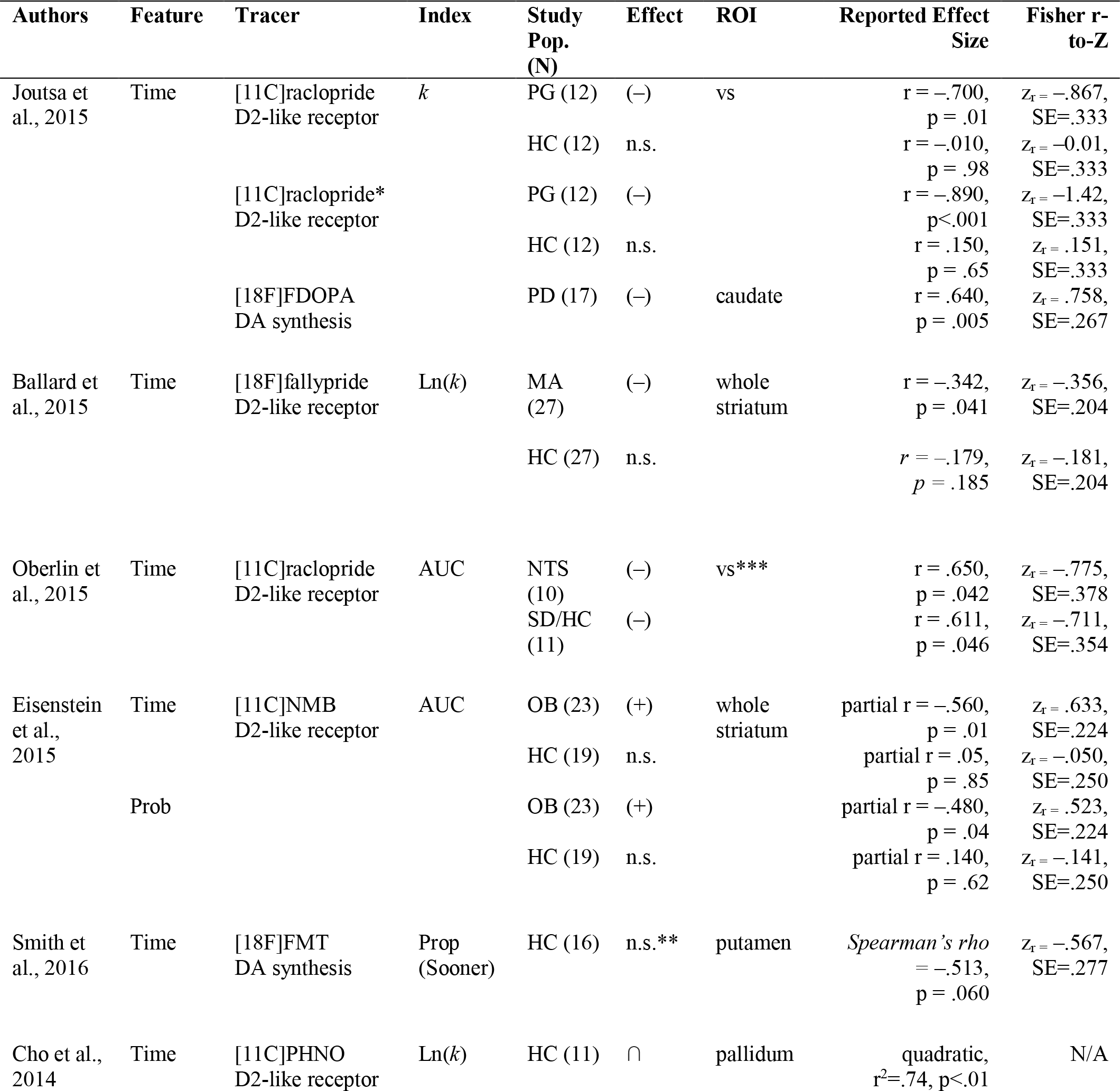
Summary of past reward discounting studies using PET imaging. Note that effect sizes are shown as originally reported but Fisher r-to-Z values have been sign-flipped when necessary to facilitate comparison of discount measures across studies (more positive values reflected greater discounting)

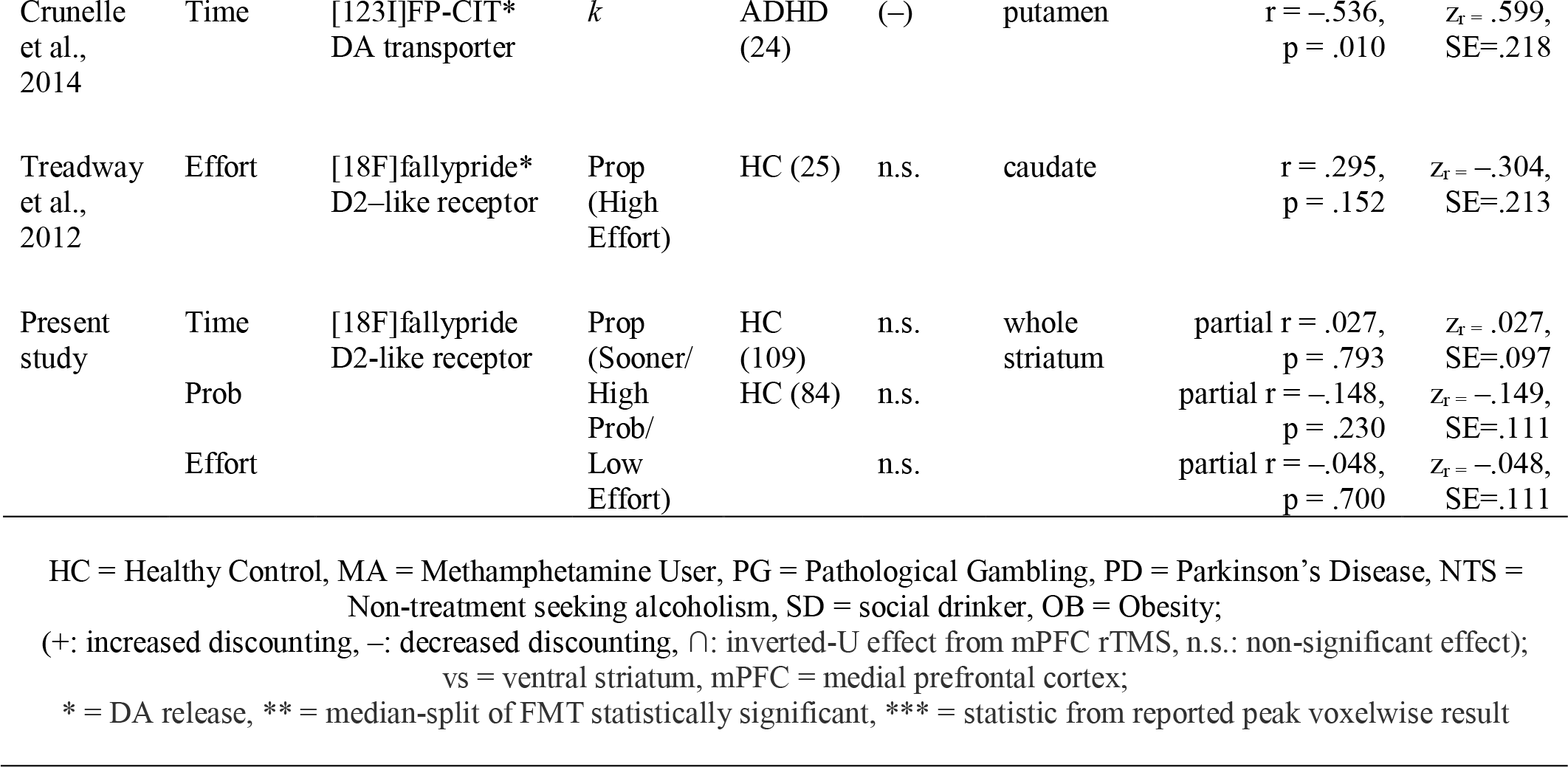

It remains unclear whether there exists a reliable association between individual differences in DA and discounting in healthy humans. Here, in two studies, one empirical and one meta-analytic, we sought to characterize the relationship between individual differences in DA and decision making in healthy human adults. In study 1, we analyzed data from three samples of healthy adults (young adults, N=25, and adult life-span, N=84, N=35). We estimated time, probability, and effort discounting of monetary rewards using multiple tasks that attempted to dissociate discounting of these three decision features and estimated DA D2-like receptor availability using PET imaging with two different radiotracers, [18F]fallypride and [11C]FLB 457, with complementary coverage of striatal and extra-striatal brain regions (Cropley et al., 2006). We analyzed data from multiple samples across a broad age range to examine the generality of effects across human adults. In study 2, we performed a quantitative meta-analysis to examine the consistency of or variation in individual differences across PET imaging studies of DA and discounting in healthy human adults and clinical groups.

## Materials and Methods

### Study 1

#### Participants and procedures

The data analyzed here were collected from three different samples at two different universities. They will be described as samples 1–3. Sample 1 included twenty-five healthy young adults (ages 18–24, M=20.9, SD=1.83, 13 females) recruited from the Vanderbilt University community in Nashville, TN between 2012 and 2013. Sample 2 included 84 healthy adults (ages 22-83, M=49.4, SD=17.6, 48 females) recruited from the Greater Nashville, TN metropolitan area between 2013 and 2016. Sample 3 included 35 healthy adults (ages 26–79, M=47.7, SD=17.4, 30 females) recruited from the Greater New Haven, CT metropolitan area between 2015 and 2017. Data from samples 1 and 2 were collected at Vanderbilt University and data from sample 3 were collected at Yale University. See Table 2 for descriptive statistics for each sample.

**Table 2.** Study demographics and decision preference descriptive statistics. Note: the difference in years of education between samples is due to Sample 1 being composed almost entirely of current college students who had not yet completed their education.

|  | Sample 1 | Sample 2 | Sample 3 |  |
| --- | --- | --- | --- | --- |
| Tracer | [18F]fallypride | [18F]fallypride | [11C]FLB 457 |  |
| N | 25 | 84 | 35 | - |
| Age | 20.9 ± 1.83 | 49.4 ± 17.6 | 47.7 ± 17.4 | $F(2,141) = 31.8, p < .001$ |
| Sex | 13 F, 12 M | 48 F, 36 M | 20 F, 15 M | $\chi^2(2, N=144) = .22, p = .896$ |
| Years Education | 14.8 ± 1.35 | 16.1 ± 1.97 | 16.5 ± 2.54 | $F(2,132) = 5.49, p = .005$ |
| Household Income | - | \$60K – 69K | \$50K – \$59K | $F(1,116) = 3.70, p = .057$ |
| Prop(sooner) | .550 ± .230 | .452 ± .243 | .497 ± .258 | $F(2,140) = 1.67, p = .191$ |
| Ln($k+1$) time | .013 ± .013 | .011 ± .013 | .014 ± .013 | $F(2,140) = .990, p = .376$ |
| Prop(high probability) | - | .681 ± .168 | .678 ± .181 | $F(1,117) = .009, p = .925$ |
| Ln($k+1$) probability | - | 1.18 ± .557 | 1.23 ± .664 | $F(1,117) = .127, p = .722$ |
| Prop(low effort) | - | .131 ± .165 | - | - |
| Ln($k+1$) effort | - | .399 ± .517 | - | - |
| Midbrain BP<sub>ND</sub> | 1.53 ± .242 | 1.20 ± .211 | 1.73 ± .407 | $F(2,141) = 48.5, p < .001$ |
| Caudate BP<sub>ND</sub> | 27.5 ± 5.27 | 25.7 ± 5.58 | - | $F(1,107) = 2.03, p = .157$ |
| Putamen BP<sub>ND</sub> | 34.1 ± 4.77 | 32.7 ± 5.09 | - | $F(1,107) = 1.48, p = .226$ |
| Ventral Striatum BP<sub>ND</sub> | 32.1 ± 8.82 | 39.2 ± 7.64 | - | $F(1,107) = 15.4, p < .001$ |
| ACC BP<sub>ND</sub> | 1.08 ± .433 | .743 ± .262 | 1.20 ± .381 | $F(2,141) = 27.9, p < .001$ |
| Thalamus BP<sub>ND</sub> | 2.63 ± .357 | 2.45 ± .409 | 3.50 ± .625 | $F(2,141) = 64.9, p < .001$ |
| Amygdala BP<sub>ND</sub> | 2.87 ± .579 | 3.18 ± .666 | 3.08 ± .798 | $F(2,141) = 1.95, p = .146$ |
| Hippocampus BP<sub>ND</sub> | 1.45 ± .703 | 1.51 ± .500 | 1.13 ± .436 | $F(2,141) = 6.47, p = .002$ |
| Insula BP<sub>ND</sub> | 2.84 ± .465 | 2.41 ± .719 | 1.86 ± .548 | $F(2,141) = 17.7, p < .001$ |

#### Screening criteria

Across samples, participants were subject to the following exclusion criteria: any history of psychiatric illness on a screening interview (a Structural Interview for Clinical DSM-IV Diagnosis was also available for all subjects and confirmed no history of major Axis I disorders) (First et al., 1997), any history of head trauma, any significant medical condition, or any condition that would interfere with MRI (e.g. inability to fit in the scanner, claustrophobia, cochlear implant, metal fragments in eyes, cardiac pacemaker, neural stimulator, pregnancy, and metallic body inclusions or other contraindicated metal implanted in the body).

Participants with major medical disorders including diabetes and/or abnormalities on a comprehensive metabolic panel, complete blood count, or EKG were excluded. Participants were also excluded if they reported a history of substance abuse, current tobacco use, alcohol consumption greater than 8 ounces of whiskey or equivalent per week, use of psychostimulants (excluding caffeine) more than twice at any time in their life or at all in the past 6 months, or any psychotropic medication in the last 6 months other than occasional use of benzodiazepines for sleep. Any illicit drug use in the last 2 months was grounds for exclusion, even in participants who did not otherwise meet criteria for substance abuse. Urine drug tests were administered, and subjects testing positive for the presence of amphetamines, cocaine, marijuana, PCP, opiates, benzodiazepines, or barbiturates were excluded. Pre-menopausal females had negative pregnancy tests at intake and on the day of the scan. There were minor differences in exclusion thresholds between samples 1/2 and sample 3 based on the location and full study protocol (e.g., a subset of subjects in sample 3 also received an oral dose of d-amphetamine). For full screening details see (Smith et al., 2017).

#### PET imaging: [18F]fallypride data acquisition and preprocessing (Samples 1 and 2)

[18F]fallypride, (S)-N-[(1-allyl-2-pyrrolidinyl)methyl]-5-(3[18F]fluoropropyl)-2,3-dimethoxybenzamide was produced in the radiochemistry laboratory attached to the PET unit at Vanderbilt University Medical Center, following synthesis and quality control procedures described in US Food and Drug Administration IND 47,245. PET data were collected on a GE Discovery STE (DSTE) PET scanner (General Electric Healthcare, Chicago, IL, USA). Serial scan acquisition was started simultaneously with a 5.0 mCi (185 MBq; study 1 median specific activity = 5.33 mCi, SD = .111; study 2 median specific activity = 5.32, SD = .264) slow bolus injection of DA D2/3 tracer [18F]fallypride (specific activity greater than 3000 Ci/mmol). CT scans were collected for attenuation correction prior to each of the three emission scans, which together lasted approximately 3.5 h with two breaks for subject comfort. Prior to the PET scan, T1-weighted magnetic resonance (MR) images (TFE SENSE protocol; Act. TR = 8.9 ms, TE = 4.6 ms, 192 TFE shots, TFE duration = 1201.9 s, FOV = 256 × 256 mm, voxel size =1 × 1 × 1 mm) were acquired on a 3T Philips Intera Achieva whole-body scanner (Philips Healthcare, Best, The Netherlands).

#### PET imaging: [11C]FLB 457 data acquisition and preprocessing (Sample 3)

[11C]FLB 457, 5-bromo-N-[[(2S)-1-ethyl-2-pyrrolidinyl]methyl]-3-methoxy-2-(methoxy-11C) benzamide was synthesized as previously described (Sandiego et al., 2015) in the radiochemistry laboratory within the Yale PET Center in the Yale School of Medicine. PET scans were acquired on the high resolution research tomograph (HRRT; Siemens Medical Solutions, Knoxville, TN, USA). [11C]FLB-457 (median specific activity: 7.80 mCi/nmol) was injected intravenously as a bolus (315 MBq; average = 8.62 mCi, SD = 2.03) over 1 min by an automated infusion pump (Harvard Apparatus, Holliston, MA, USA). Prior to each scan, a six-minute transmission scan was performed for attenuation correction. Dynamic scan data were acquired in list mode for 90 min following the administration of [11C]FLB 457 and reconstructed into 27 frames (6 × 0.5 mins, 3 × 1 min, 2 × 2 mins, 16 × 5 mins) with corrections for attenuation, normalization, scatter, randoms, and dead time using the MOLAR (Motion-compensation OSEM List-mode Algorithm for Resolution-Recovery Reconstruction) algorithm (Carson et al., 2004). Event-by-event, motion correction (Jin et al., 2013) was applied using a Polaris Vicra optical tracking system (NDI Systems, Waterloo, Canada) that detects motion using reflectors mounted on a cap worn by the subject throughout the duration of the scan. Prior to the PET scan, T1-weighted magnetic resonance (MR) images (MPRAGE protocol; TR = 2.4 s, TE = 1.9 ms, FOV = 256 × 256 mm, voxel size = 1 × 1 × 1 mm) were acquired on a 3T Trio whole-body scanner (Siemens Medical Systems, Erlangen, Germany). After decay correction and attenuation correction, PET scan frames were corrected for motion using SPM8 (Friston et al., 1994) with the 20th dynamic image frame of the first series serving as the reference image. The realigned PET frames were then merged and re-associated with their acquisition timing info in PMOD’s PVIEW module to create a single 4D file for use in PMOD’s PNEURO tool for further analysis.

#### Binding Potential Calculation

We estimated D2 receptor availability as binding potential (BP_ND_) using the simplified reference tissue model (SRTM) with the cerebellum as the reference region) (Lammertsma and Hume, 1996) via two approaches: voxelwise and ROI-based (by fitting time activity curves). PMOD’s PXMOD tool was used to estimate BP_ND_ voxel-wise using a published basis function fitting approach (Gunn et al., 1997). See Figure 1 for average voxelwise BP_ND_ images from all three samples.

**Figure 1.**
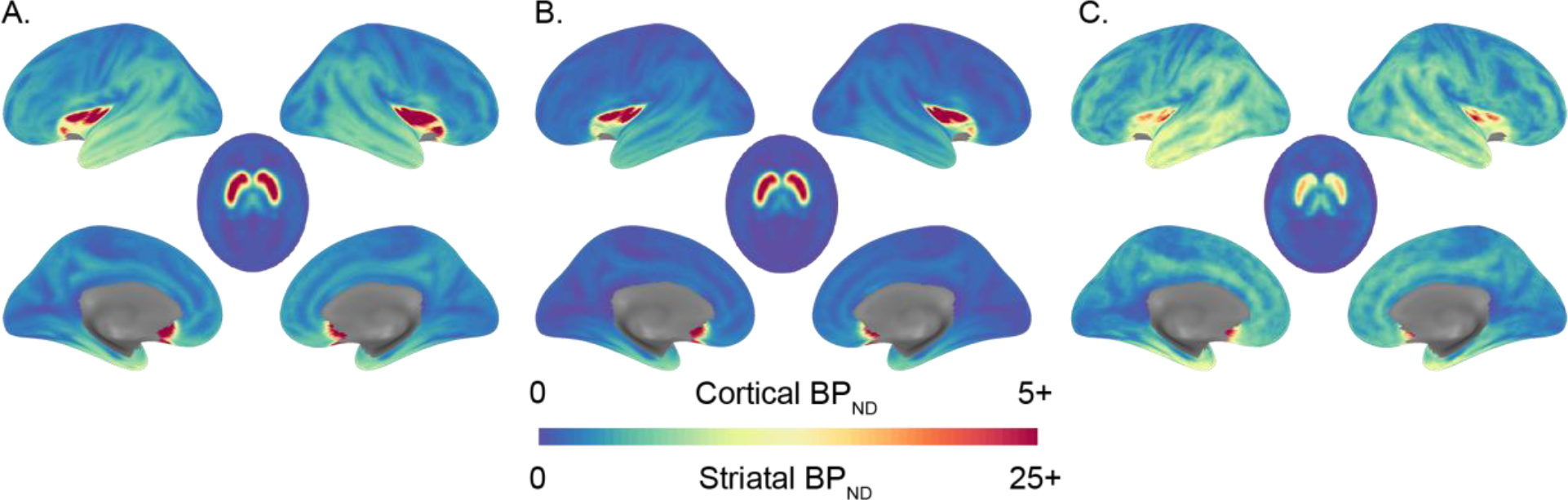
Average dopamine D2-like receptor availability. Average voxelwise whole-brain binding potential for (A) Sample 1 collected using [18F]fallypride in young adults, (B) Sample 2 collected using [18F]fallypride across the adult life span, and (C) Sample 3 collected using [11C]FLB 457 across the adult life span. Sagittal images use the cortical BP_ND_ color scale and axial images use the striatal BP_ND_ color scale. Note the differences in binding potential between cortical and striatal regions depend on the radiotracer and mean age of the sample.

The set of regions of interest did not completely overlap across samples due to differences in regional coverage of the radiotracers (samples 1, 2, 3: midbrain, thalamus, amygdala, hippocampus, anterior cingulate cortex (ACC), and insula; samples 1 and 2: ventral striatum, caudate, putamen). The midbrain was drawn in MNI standard space using previously described guidelines (Mawlawi et al., 2001; Dang et al., 2012b; Dang et al., 2012a) and registered to PET images using the same transformations used in BP_ND_ calculation. All other ROIs were derived from the Hammers Atlas plus deep nuclei parcellation as produced from the parcellation of the T1 structural image of each subject in the PNEURO module of PMOD software. The PET data was registered to the T1 image for each subject and, thus, to the ROIs (all steps implemented in PNEURO module of PMOD Software). BP_ND_ values from ROIs were obtained by fitting the SRTM to the PET time activity curve data from each ROI in the PKIN (kinetic modeling) module of PMOD using the cerebellum as the reference region. These ROI-based BP_ND_ values were then averaged across hemispheres. Recently, our lab and others have shown that many brain regions may be susceptible to partial volume effects in estimating BP_ND_ especially in older adults as a result of age differences in gray matter volume (Smith et al., 2017). PVC increased estimated binding potential across adults of all ages while also increasing individual differences not related to age (Smith et al., 2017). Therefore, we used PVC values in all analyses presented here with the exception of the midbrain for which we used uncorrected BP_ND_ for analysis, because it was not available in the Hammers Atlas in PNEURO. We shared both corrected and uncorrected values for all ROIs if others want to do additional analysis. These data can be accessed at https://osf.io/htq56/.

We extracted mean D2-like BP_ND_ from the midbrain (mean ± SD:1.39 ± .356) for all samples since both [18F]fallypride and [11C]FLB 457 have demonstrated good signal-to-noise ratio (SNR) in this region (Ray et al., 2012; Narendran et al., 2014). We extracted mean striatal D2-like BP_ND_ from samples 1 and 2 in the ventral striatum (uncorrected mean ± SD: 18.6 ± 3.30; PVC mean ± SD: 37.6 ± 8.43), caudate (uncorrected mean ± SD: 16.18 ± 3.52; PVC mean ± SD: 26.1 ± 5.54), and putamen (uncorrected mean ± SD: 22.8 ± 3.40; PVC mean ± SD: 33.0 ± 5.03).

Since [11C]FLB 457 has poor SNR in the striatum compared to [18F]fallypride, we did not extract striatal BP_ND_ from sample 3. We extracted mean D2-like BP_ND_ from all samples in the anterior cingulate cortex (ACC) (uncorrected mean ± SD: .732 ± .281; PVC mean ± SD: .912 ± .385), thalamus (uncorrected mean ± SD: 2.32 ± .622; PVC mean ± SD: 2.74 ± .638), amygdala (uncorrected mean ± SD: 2.191 ± .490; PVC mean ± SD: 3.10 ± .692), hippocampus (uncorrected mean ± SD: 1.05 ± .308; PVC mean ± SD: 1.40 ± .546), and insula (uncorrected mean ± SD: 2.12 ± .654; PVC mean ± SD: 2.35 ± .714). To avoid arbitrary delineations of larger cortical regions, cortical BP_ND_ associations were evaluated with whole-brain voxelwise analyses (discussed in *Experimental Design and Statistical Analysis*).

Approval for the [18F]fallypride study protocol (samples 1 and 2) was obtained from the Vanderbilt University Human Research Protection Program and the Radioactive Drug Research Committee. Approval for the [11C]FLB 457 study protocol (sample 3) was obtained from the Yale University Human Investigation Committee and the Yale-New Haven Hospital Radiation Safety Committee. All participants in each sample completed written informed consent. Each samples’ study procedures were approved in accordance with the Declaration of Helsinki’s guidelines for the ethical treatment of human participants.

#### Reward discounting tasks

All samples completed a temporal discounting task (N=144), samples 2 and 3 also completed a probability discounting task (N=119), and sample 2 also completed a physical effort discounting task (N=84). All tasks were incentive-compatible (played for real cash earnings) and performed during fMRI scanning (samples 1 and 2) or on a computer in a behavioral lab (sample 3) on a separate visit from the PET imaging session as part of larger multimodal neuroimaging studies. The average number of days between a PET imaging session and performance on discounting tasks was similar between studies (sample 1: 18.2±12.5, sample 2: 25.0±18.4, sample 3: 38.9±27.3).

#### Temporal discounting task

All three samples completed a temporal discounting task adapted from a previously used paradigm (McClure et al., 2004). On each trial, participants chose between an early monetary reward and a late reward. In sample 1, the delay of the early reward was set to today, 2 weeks, or 1 month, while the delay of the late reward was set to 2 weeks, 1 month, or 6 weeks. In samples 2 and 3, the delay of the early reward was set to today, 2 months, or 4 months, while the delay of the late reward was set to 2 months, 4 months, or 6 months. In all samples, the early reward magnitude ranged between 1% and 50% less than the late reward. Participants in sample 1 played 84 (42 trials in two runs) trials of the temporal discounting task and participants in samples 2 and 3 played 82 trials (41 trials in two runs). One participant in sample 3 had missing data for this task, producing a total sample size of 143 participants with temporal discounting data across all samples.

#### Probabilistic discounting task

Samples 2 and 3 completed a probabilistic decision making task similar to commonly used two-alternative forced choice mixed gamble tasks (Levy and Glimcher, 2011). On each trial, participants chose between a smaller monetary reward with a higher probability and a larger reward with a lower probability. The probability of the higher probability reward was set to 50%, 75%, or 100%, while the probability of the lower probability reward was set to 25% or 50% lower. The higher probability reward magnitude ranged between 1% and 50% lower compared to the lower probability reward. Participants in samples 2 and 3 played 82 trials of the probability discounting task. Data for this task was available for all participants, producing a total sample size of 119 participants with probability discounting data.

#### Effort discounting task

The Effort Expenditure for Rewards Task (EEfRT) was adapted from an existing paradigm that used finger pressing as the physical effort required for earning a reward (Treadway et al., 2009). On each trial, participants chose between a smaller monetary reward available for a lower amount of physical effort (pinky finger button presses) and a larger reward available for a higher amount of effort. The effort required for the smaller reward was set as 35%, 55%, or 75% (of each participant’s maximum press rate), while the effort required for the larger reward was set as 20% or 40% higher than the smaller reward (i.e., 55%, 75%, or 95%). The number of button presses required for each level of effort was individually determined based on an initial calibration procedure in which participants pressed a button with their pinky finger as many times and as rapidly as possible in a few short intervals. The smaller magnitude reward ranged between 1% and 50% lower than the larger reward. On half of the trials, after making a choice participants were shown a 1-second “Ready” screen and then completed the button-pressing task. Participants in sample 2 played 82 trials of the effort discounting task. No participant had missing data for this task, producing a total sample size of 84 participants with effort discounting data.

#### Computational modeling of reward discounting

In addition to a simple calculation of the proportion of smaller magnitude (less delayed/higher probability/lower effort) reward choices, we used a computational model to estimate behavioral preferences. For each participant and each task, discounting was modeled with a hyperbolic discounted value function,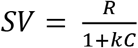,where *R* represents the monetary reward magnitude, *k* represents the discount rate, and *C* represents either: (1) proportion of maximum finger press rate for effort, (2) odds against winning (1-*p*(win))/*p*(win)) for probability, or (3) delay in days for time. Data were fit with a softmax as the slope of the decision function. Since *k* values are not normally-distributed, we used natural log-transformed values Ln(*k*+1). Past work from our lab has shown *k* values and simple proportion of smaller reward choices are highly correlated (Seaman et al., 2018). We report both scores for transparency.

#### Experimental Design and Statistical Analysis

To determine whether D2-like receptor availability in the midbrain, striatum, and extrastriatal regions were associated with discounting, we combined one sample of healthy young adults with two cross-sectional healthy adult life-span samples. We ran linear regressions between BP_ND_ and the proportion of sooner/higher probability/lower effort choices as well as *k*-values. Regressions included control variables for age, sex, and study sample (using dummy coded variables for samples 2 and 3 where appropriate). Standardized beta coefficients are reported for these primary analyses. We corrected for multiple comparisons within each cost domain (time, probability, effort) for each region available for each combination of samples since not all samples were tested on all tasks or had BP_ND_ for all regions. We applied Bonferroni-correction to *p*-values as follows: midbrain =.05; striatal ROIs = .05/3 =.016; extrastriatal ROIs = .05/5 =.010. Previous work has documented associations between discounting and household income and education (de Wit et al., 2007; Reimers et al., 2009). Since we did not identify such associations between education or income with discounting in any task, we did not include these measures as covariates in regressions.

Additional exploratory ROI analyses examined whether associations between dopamine and discounting varied across age groups or study samples. Full evaluation of these effects required running 27 additional multiple regression analyses that evaluated main effects of D2-like receptor availability, sex, age, and study sample (as above in the primary analyses) in addition to interactions between age and D2-like receptor availability and study sample and D2-like receptor availability. Given the lack of specific hypotheses for these exploratory analyses, we applied a Bonferroni correction for multiple comparisons; only interactions that were significant at *p* < .00185 (i.e., .05/27 = .00185) are reported with follow-up within-group tests. Interactions are reported as unstandardized beta coefficients. Full model outputs for all of these analyses are available on OSF: https://osf.io/htq56/.

Exploratory voxelwise statistical testing of D2-like receptor availability was separately carried out for each discounting task in each sample in MNI standard space. Since [11C]FLB 457 was acquired on a high resolution scanner which produced maps with lower local spatial correlation, we spatially smoothed these BP_ND_ maps with a 5mm FWHM Gaussian kernel to increase spatial SNR (Christopher et al., 2014; Plaven-Sigray et al., 2017). Linear regressions examining the effect of proportion of sooner/higher probability/lower effort choices or Ln(*k*+1) values on voxelwise BP_ND_ with age and sex as covariates were carried out using FSL Randomise (Version 2.9) within each sample. Threshold-free cluster enhancement (Smith and Nichols, 2009) was used to detect regions with significant correlations across the whole brain with non-parametric permutation tests (5,000 permutations). Statistical maps were thresholded at *p* < 0.05.

### Study 2

To identify research studies of interest, a PubMed search for the following terms (((Dopamine) AND positron emission tomography) AND humans) AND (discounting OR impulsive choice) yielded 10 studies. Five of these studies included original analysis of the relationship between preferences in a discounting task and a PET measure of DA function and were included. An additional exhaustive search via Google Scholar identified 3 additional relevant and includable studies. Notably, six of the studies in the meta-analysis used tracers that bind to D2-like receptors for baseline receptor availability or DA release measures (Treadway et al., 2012; Ballard et al., 2015; Cho et al., 2015; Eisenstein et al., 2015; Joutsa et al., 2015; Oberlin et al., 2015), two used tracers that measure presynaptic DA uptake (Joutsa et al., 2015; Smith et al., 2016), and one used a tracer that binds to dopamine transporters (DAT) (Crunelle et al., 2014). The study measuring DAT reported methylphenidate (MPH) occupancy after drug administration. To obtain the DAT BP_ND_ measure, we sign-flipped the correlation since DAT BP_ND_ is inversely related to MPH occupancy. In addition to the present study (Study 1) that examined time, probability, and effort, one other study examined both time and probability discounting (Eisenstein et al., 2015), another study examined effort-based discounting (Treadway et al., 2012), and the remaining studies examined only time discounting. One of these studies used single photon emission computerized tomography (SPECT) rather than PET and was included. If correlation coefficients were not reported, t-statistics and degrees of freedom were used to generate correlation coefficients using the formula 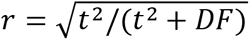 (Rosenthal and Rosnow, 2008). Because correlations are bound and can be skewed, they were Fisher r-to-Z transformed before meta-analysis. In the case of one study (Treadway et al., 2012), correlations between caudate D2-like receptor BP_ND_ and preferences for effort were originally reported as three within-task correlations (by probability condition). To approximate the full task correlation, we used the Fisher r-to-Z transformation for the three correlations and then averaged these values. Depending on the decision preference index reported (Ln(*k*), proportion smaller, area-under-the-curve, etc.), we sign-flipped Z-scores so that more positive values reflected greater discounting (e.g., less willing to choose a larger, delayed/uncertain/effortful reward). One study did not assess or report linear correlations (Cho et al., 2015). A summary of these studies is presented in Table 1.

Since our group previously reported little to no correlation between time, probability, and effort discounting in sample 2 (Seaman et al., 2016), we limited the meta-analysis to time discounting measures only. Therefore, the meta-analysis included 7 studies with 14 correlation effects (including the effect of time discounting from the present study). The goal of the meta-analysis was to identify generalizable patterns that address the broader question of whether discounting is related to general striatal dopamine function. Since prior reports indicated positive associations within individuals between tracer targets (e.g., D2 receptors and DAT (Volkow et al., 1998; Yang et al., 2004), D2 receptors and DA synthesis capacity (Berry et al., 2018), D2 receptors and DA release (Samanez-Larkin et al., 2013), DAT and DA synthesis capacity (Sun et al., 2012), DAT and DA release (Volkow et al., 2002)), we included all studies that reported a correlation with a striatal region. It should be noted that indices of any one of these radiotracer targets alone may not be reflective of general dopamine function, but contribute to and interact within complex spatiotemporal circuits that impact dopaminergic synapses. If a study reported multiple striatal regions, we used the reported t-statistics and p-values to select only the region with the largest effect size. Since this resulted in inconsistent ROIs (with 6 effects in the whole striatum, 6 in the ventral striatum, 2 in the caudate, and 2 in the putamen), we compared the correlation between time discounting in the present study with D2-like receptor availability in the whole striatum. BP_ND_ for the whole striatum was calculated as a volume-weighted average of the caudate, putamen, and ventral striatum PVC BP_ND_ values. Included effects from the present study controlled for age, sex, and study sample. Replacing the whole striatum value with the largest substriatal effect size value in our study (ventral striatum) did not change the pattern of results.

Meta-analytic effects were derived using the metafor R package (Viechtbauer, 2010) in JASP (Version 0.8.5.1) using random effects with restricted maximum-likelihood (JASP Team, 2018) to help account for between-study variance. An initial meta-analysis across all studies evaluated whether the common correlation (intercept) was significantly greater than zero, *p* < .05. Since the study samples included groups with psychopathology and radiotracers that bind to different dopaminergic targets, we ran additional meta-analytic models to evaluate whether effect sizes depended on the interaction of these terms. We dummy-coded study populations as either belonging to a group that is characterized by addiction, healthy controls, or any other psychopathology or disease. We coded the following as addiction: pathological gambling, methamphetamine users, and non-treatment-seeking alcoholism. Other psychopathology samples included obesity, PD, and treatment-naïve ADHD samples. Radiotracer targets were either D2-like receptors (D2R) including baseline and release measures, DA synthesis capacity (SC), or dopamine transporters (DAT). We used the Q-statistic to test the null hypothesis that the common true correlation is zero and *I*^*2*^ values to assess significance due to variance explained by heterogeneity of the effects (Borenstein et al., 2011). Model fit quality statistics are reported for the intercept model and the interaction model, along with each of the interaction main effect terms alone. We evaluated publication bias and study precision asymmetry with visual inspection of a funnel plot and Egger’s test (*p* < .05).

## Results

### Study 1

#### Discounting across studies

Average behavioral measures of time and probability discounting did not differ between samples (time: *F*(2,140) = 1.63, *p* = .200; probability: *F*(1,117) = .009, *p* = .925), facilitating our ability to combine samples for analysis. Simple choice proportions (e.g., smaller-sooner / total number of choices) were highly correlated with computationally-estimated discount rates Ln(*k*+1) for time (r_141_ = .829, *p* < .001), probability (r_117_ = .798, *p* < .001), and effort (r_82_ = .830, *p* < .001). A previous publication documented a lack of associations between time, probability, and effort discounting within a subset of sample 2 (N=75) with the exception of a modest significant correlation between time and effort discounting using the proportion choice variable but not using the discounting parameters from the hyperbolic models (Seaman et al., 2018). Across the samples included here, we also observed a general lack of associations between discounting across the tasks. Once again, the only exceptions were significant correlations within sample 2 between time and effort discounting using both the proportion choice variables (r_82_ = .27, p = .014) and, here, a significant correlation between the proportion choice variable for time discounting and the discounting model parameter (Ln(*k*+1)) for effort discounting (r_82_ = .27, p = .012). However, note that any associations or lack of associations with behavioral measures of effort discounting should be viewed with caution given that most participants selected a high proportion of larger/high-effort choices creating a ceiling effect that restricted the range of values.

#### Age effects on Discounting and D2-like receptor availability

Samples 2 and 3 included adults of all ages. Age was not reliably associated with reward discounting of time (r_141_ = .049, *p* = .563), probability (r_117_ = −.007, *p* = .947), or effort (r_82_ = .116, *p* = .293). Age was negatively correlated with BP_ND_ in the midbrain (r_142_ = −.442, 95% CI [−.565, −.300], *p* < .001), caudate (r_107_ = −.409, 95% CI [−.555, −.240], *p* < .001), putamen (r_107_ = −.350, 95% CI [−.505, −.173], *p* < .001), anterior cingulate (r_142_ = −.316, 95% CI [−.456, −.161], *p* < .001), and insula (r_142_ = −.437, 95% CI [−.560, −.294], *p* < .001) but not in the ventral striatum (r_107_ = .083, 95% CI [−.106, − .267], *p* = .389), amygdala (r_142_ = −.145, 95% CI [−.301, .019], *p* = .083), hippocampus (r_142_ = − .130, 95% CI [−.287, .034], *p* = .121), or thalamus (r_142_ = −.125, 95% CI [−.283, .039], *p* = .136).

Correlations between age and discounting within sample 2 were previously reported in (Seaman et al., 2018). Correlations between age and BP_ND_ for samples 2 and 3 were previously reported in (Dang et al., 2016) and (Smith et al., 2017).

#### Discounting and D2-like receptor availability

We did not identify associations between D2-like BP_ND_ in the midbrain and discounting across samples 1, 2, and 3 or the striatum and discounting across samples 1 and 2 (Table 3). We identified a modest positive correlation between probability discounting and D2-like receptor availability in the hippocampus (Ln(*k*+1): β = .197, SE = .110, t_114_ = 2.06, *p* = .042). However, the correlation did not survive correction for multiple comparisons. No associations were identified between discounting and any of the other ROIs in the primary analyses (Table 3 and Figure 2).

**Table 3.**
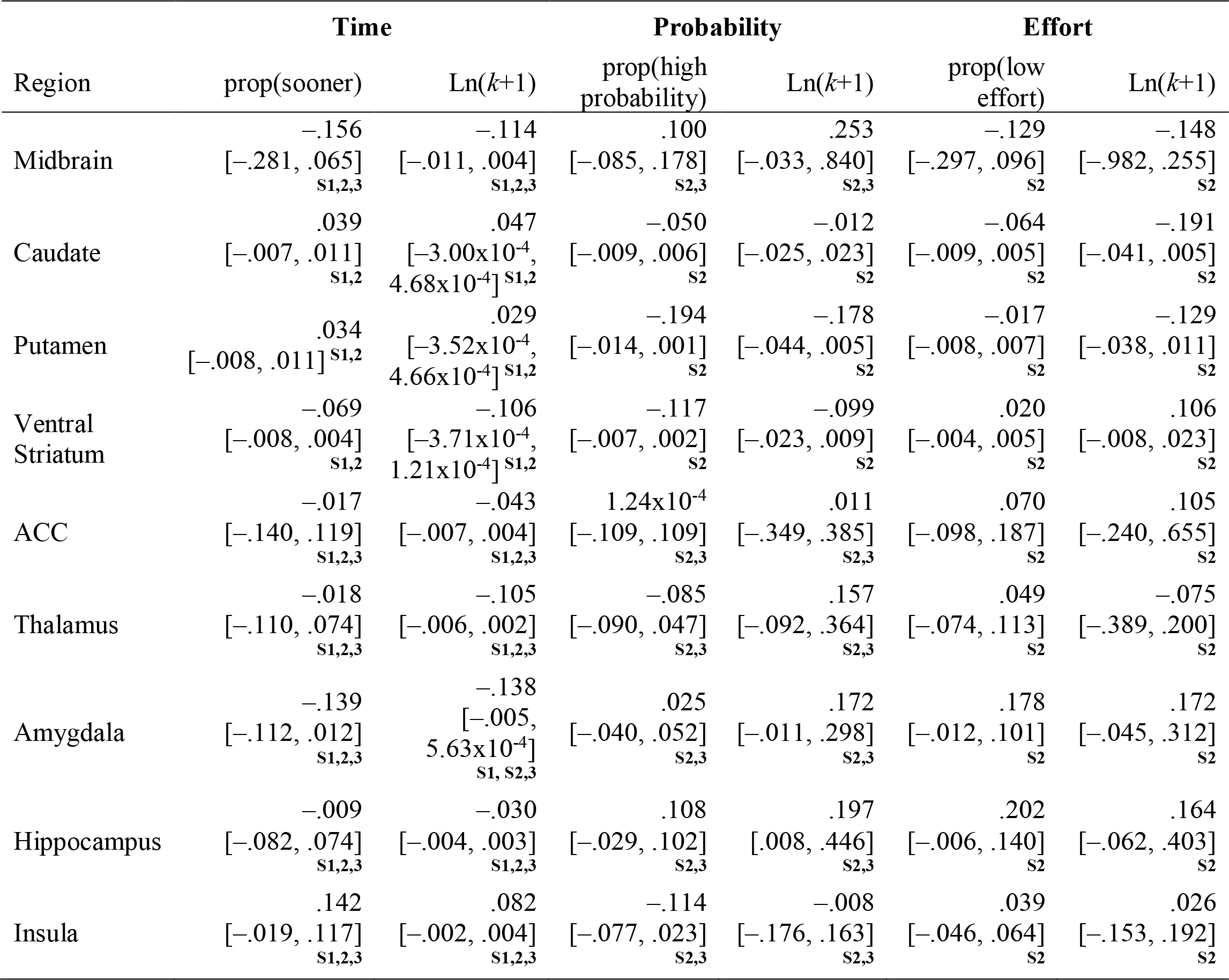
Region of interest analyses for D2-like receptor availability (PVC) showing standardized regression coefficients (after adjustment for control variables) and 95% confidence intervals. S1 = sample 1, S2 = sample 2, S3 = sample 3.

| Region | Time |  | Probability |  | Effort |  |
| --- | --- | --- | --- | --- | --- | --- |
|  | prop(sooner) | Ln(*k*+1) | prop(high probability) | Ln(*k*+1) | prop(low effort) | Ln(*k*+1) |
| Midbrain | -.156<br>[-.281, .065]<br>S1,2,3 | -.114<br>[-.011, .004]<br>S1,2,3 | .100<br>[-.085, .178]<br>S2,3 | .253<br>[-.033, .840]<br>S2,3 | -.129<br>[-.297, .096]<br>S2 | -.148<br>[-.982, .255]<br>S2 |
| Caudate | .039<br>[-.007, .011]<br>S1,2 | .047<br>[-3.00x10<sup>-4</sup>, 4.68x10<sup>-4</sup>]<br>S1,2 | -.050<br>[-.009, .006]<br>S2 | -.012<br>[-.025, .023]<br>S2 | -.064<br>[-.009, .005]<br>S2 | -.191<br>[-.041, .005]<br>S2 |
| Putamen | .034<br>[-.008, .011]<br>S1,2 | .029<br>[-3.52x10<sup>-4</sup>, 4.66x10<sup>-4</sup>]<br>S1,2 | -.194<br>[-.014, .001]<br>S2 | -.178<br>[-.044, .005]<br>S2 | -.017<br>[-.008, .007]<br>S2 | -.129<br>[-.038, .011]<br>S2 |
| Ventral Striatum | -.069<br>[-.008, .004]<br>S1,2 | -.106<br>[-3.71x10<sup>-4</sup>, 1.21x10<sup>-4</sup>]<br>S1,2 | -.117<br>[-.007, .002]<br>S2 | -.099<br>[-.023, .009]<br>S2 | .020<br>[-.004, .005]<br>S2 | .106<br>[-.008, .023]<br>S2 |
| ACC | -.017<br>[-.140, .119]<br>S1,2,3 | -.043<br>[-.007, .004]<br>S1,2,3 | 1.24x10<sup>-4</sup><br>[-.109, .109]<br>S2,3 | .011<br>[-.349, .385]<br>S2,3 | .070<br>[-.098, .187]<br>S2 | .105<br>[-.240, .655]<br>S2 |
| Thalamus | -.018<br>[-.110, .074]<br>S1,2,3 | -.105<br>[-.006, .002]<br>S1,2,3 | -.085<br>[-.090, .047]<br>S2,3 | .157<br>[-.092, .364]<br>S2,3 | .049<br>[-.074, .113]<br>S2 | -.075<br>[-.389, .200]<br>S2 |
| Amygdala | -.139<br>[-.112, .012]<br>S1,2,3 | -.138<br>[-.005, 5.63x10<sup>-4</sup>]<br>S1, S2,3 | .025<br>[-.040, .052]<br>S2,3 | .172<br>[-.011, .298]<br>S2,3 | .178<br>[-.012, .101]<br>S2 | .172<br>[-.045, .312]<br>S2 |
| Hippocampus | -.009<br>[-.082, .074]<br>S1,2,3 | -.030<br>[-.004, .003]<br>S1,2,3 | .108<br>[-.029, .102]<br>S2,3 | .197<br>[.008, .446]<br>S2,3 | .202<br>[-.006, .140]<br>S2 | .164<br>[-.062, .403]<br>S2 |
| Insula | .142<br>[-.019, .117]<br>S1,2,3 | .082<br>[-.002, .004]<br>S1,2,3 | -.114<br>[-.077, .023]<br>S2,3 | -.008<br>[-.176, .163]<br>S2,3 | .039<br>[-.046, .064]<br>S2 | .026<br>[-.153, .192]<br>S2 |

**Figure 2.**
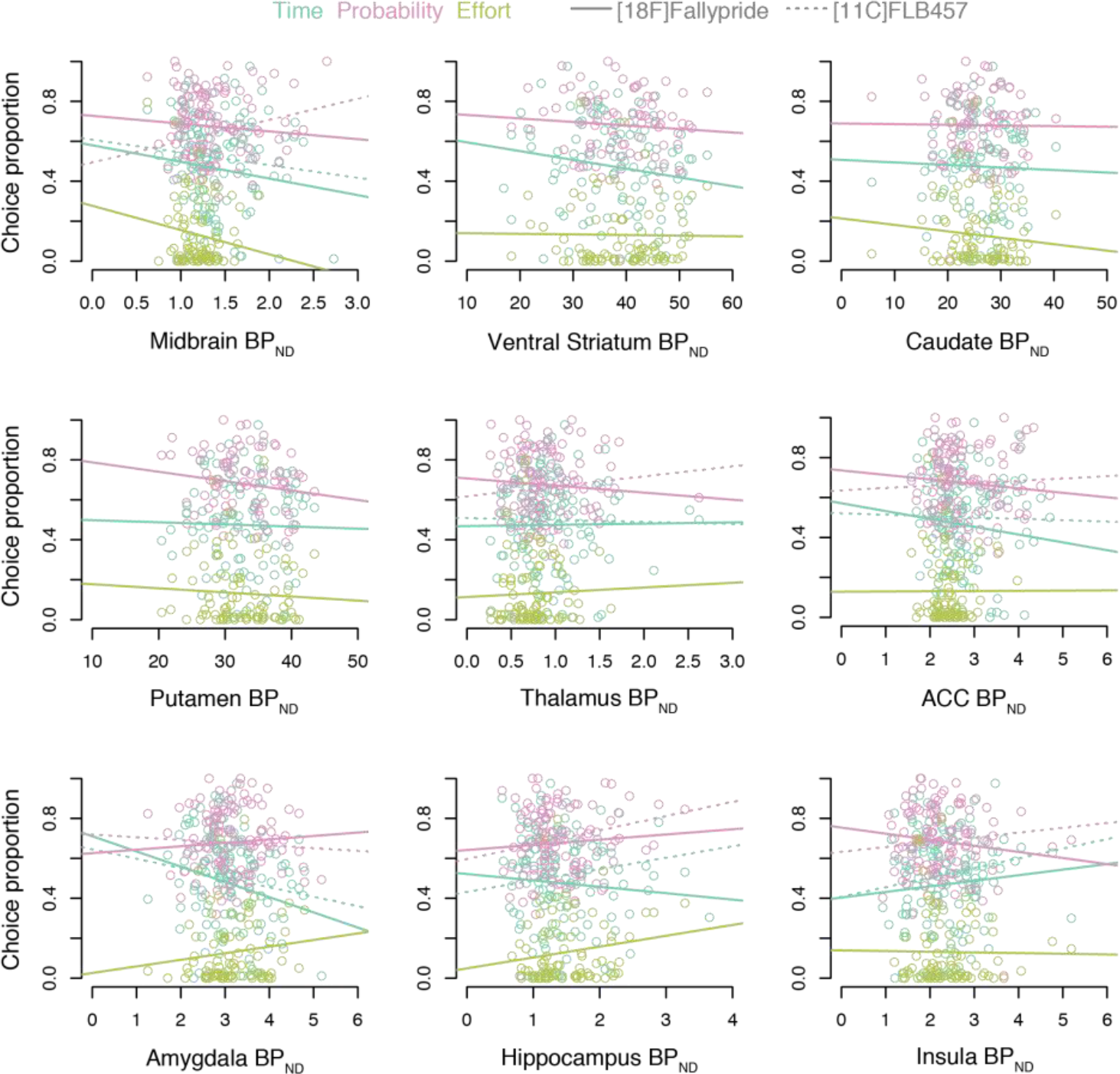
Correlations between reward discounting and D2-like receptor availability. Correlation plots depict associations between D2-like receptor availability (PVC) and proportion of smaller sooner / higher probability / less effortful choices. Individual subject data points are depicted for time in turquoise, probability in pink, and effort in green. Solid lines represent regression slopes for [18F]fallypride and dotted lines represent regression slopes for [11C]FLB 457.

Exploratory evaluation of possible interactions between age and D2-like receptor availability or study sample and D2-like receptor availability in predicting discounting revealed four significant interactions after controlling for multiple comparisons. The significant interactions revealed that the association between D2-like receptor availability and effort discounting varied across age in the midbrain (chose low effort: β = −.0195, *p* = .00006, Ln(*k*+1): β = −.0496, *p* = .0015) and ventral striatum (chose low effort: β = −.00005, *p* = .0009, Ln(*k*+1): β = −.002, *p* = .0002). Follow-up analyses examined simple effects within younger adults (ages 18-30), middle-aged adults (ages 31-57), and older adults (ages 57-83) from a tertile split of ages. In the midbrain, there was a negative association between D2-like receptor availability and discounting within older adults (chose low effort: r_32_ = −.50, *p* = .002, Ln(*k*+1): r_32_ = −.46, *p* = .006) but non-significant associations in younger adults (chose low effort: r13 = .23, *p* = .41, Ln(*k*+1): r_13_ = .25, *p* = .36) and middle-aged adults (chose low effort: r_33_ = .27, *p* = .12, Ln(*k*+1): r_33_ = .12, *p* = .48). In the ventral striatum, there was a positive association between D2-like receptor availability and discounting within younger adults (chose low effort: r_13_ = .67, *p* = .007, Ln(*k*+1): r_13_ = .76, *p* < .001) but non-significant associations in middle-aged adults (chose low effort: r_33_ = .10, *p* = .57, Ln(*k*+1): r_33_ = .20, *p* = .26), and older adults (chose low effort: r_32_ = .18, *p* = .31, Ln(*k*+1): r_32_ = −.16, *p* = .38). No age by D2-like receptor availability interactions reached corrected levels of significance in any other ROI for effort discounting and in any ROI for time and probability discounting. No study sample by D2-like receptor availability interactions reached corrected levels of significance in any ROI for any task. See OSF for complete model output and figures: https://osf.io/htq56/.

Voxelwise analysis of binding potential maps did not reveal any significant correlations with discounting. Unthresholded statistical maps can be viewed/downloaded from NeuroVault at: https://neurovault.org/collections/ZPFBVXPK/

### Study 2

#### Meta-analysis: DA PET studies of reward discounting

An initial meta-analysis across all studies of temporal discounting did not identify a significant common correlation between discounting and kinetic measure of DA function (Omnibus test of model coefficients, Cochran’s Q = 1.03, *p* = .310, *I*^2^ = 84.7%; β_intercept_ = −.167, SE = .164, Z = −1.02, AIC = 28.6).

Alternatively, a model that included the interaction between psychopathology group and radiotracer target provided a better fit than the common correlation model (without interaction terms) and accounted for the heterogeneity of effects (Omnibus test of model coefficients, Cochran’s Q = 35.2, *p* < .001, *I*^2^ = 37.15%, AIC = 19.8). Inspection of the coefficients suggested that psychopathology alone had a greater impact on the model than radiotracer target: β_Healthy,D2-receptor/intercept_ = −.088, SE = .124, Z = −.708, *p* = .479, β_DA synthesis capacity_ = −.479, SE = .357, Z = − 1.34, *p* = .180, β_DAT_ = −.034, SE = .410, Z = −.084, *p* = .933, β_Addiction_ = −.676, SE = .215, Z =−3.14, *p* = .002, β_Other Psychopathology_ = .720, SE = .317, Z = 2.27, *p* = .023, β_Other Psychopathology, DA synthesis capacity_ = .605, SE = .566, Z = 1.07, *p* = .285.

A follow-up model with the radiotracer target interaction term alone provided a worse fit (Omnibus test of model coefficients, Cochran’s Q = 2.59, *p* = .273, *I*^2^ = 83.9%, AIC = 28.4).

However, the follow-up model with the psychopathology term alone provided the best model fit compared to all other meta-analysis models (Omnibus test of model coefficients, Cochran’s Q = 35.7, *p* < .001, *I*^2^ = 31.8%, AIC = 14.3). Again, inspection of the coefficients suggested that psychopathology alone had a greater impact on the model, regardless of radiotracer target: β_Healthy/intercept_ = −.138, SE = .110, Z = −1.26, *p* = .207, β_Addiction_ = -.616, SE = .202, Z = −3.05, *p* = .002, β β_Other Psychopathology_ = .793, SE = .199, Z = 3.99, *p* < .001. A forest plot of the psychopathology model is provided in Figure 3. Plotted values depict Pearson correlation coefficients for display purposes only. Visual inspection of asymmetry in a funnel plot of effects from the psychopathology model (Figure 2) and Egger’s test (Z = −2.24, *p* = .025) indicated some potential publication bias associated with differences between studies reporting effects in specific psychopathology groups. Egger’s test did not indicate the presence of publication bias in the common correlation model (Z = −1.80, *p* = .072) or full interaction model (Z = −1.56, *p* = .119).

**Figure 3.**
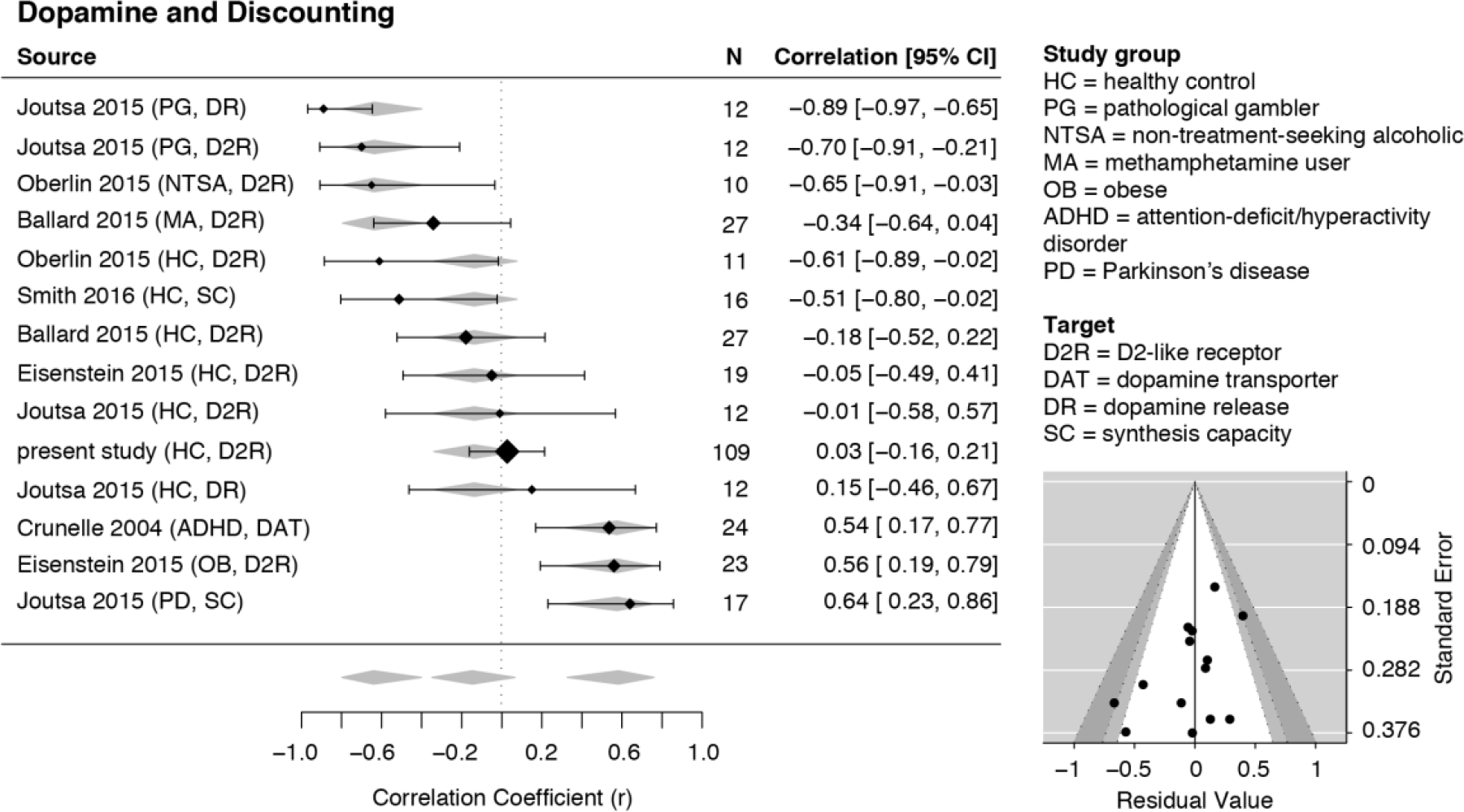
Meta-analytic comparison of associations between individual differences in dopamine and reward discounting. The forest plot on the left shows variation in effect sizes according to clinical status (healthy, addiction, and other psychopathology). Values depict correlation coefficients, r, for display purposes; positive values indicate a positive correlation between DA function and greater discounting (e.g., more immediate choices). Black diamonds represent individual study effects (diamond size depicts the weight in the meta-analysis and the horizontal lines represent 95% confidence intervals of the individual effects, noted on the right). Gray diamonds represent 95% confidence intervals of the factor coefficients from the clinical status term. The funnel plot is displayed in the lower right for the model. Plotted points represent individual effects. Points represent the residuals of the psychopathology groups and their associated study precision (standard error). When the effect residuals lie within the unshaded area, it implies that heterogeneity in the main effect is successfully accounted for by the interaction model. Points within the unshaded region correspond to p-values greater than .10 while p-values in the light gray and dark gray regions correspond to p-values between .10 and .05 and between .05 and .01, respectfully.

The nature of the psychopathology group effect was that healthy individuals showed a non-significant, small, negative correlation between DA and discounting, the addiction groups showed a significant and stronger negative association, and the other psychopathology groups showed a stronger positive association relative to healthy controls. To facilitate comparison of group effects with past and future studies, we converted estimated coefficient Z-values back to Pearson correlation coefficients. The correlation for the healthy group was r = −.137, 95% CI [−.339, .076], the correlation for the addiction group was r = −.638, 95% CI [−.796, −.399], and the correlation for the other psychopathology group was r = .575, 95% CI [.319, .753]. Including additional data from studies using effort and probability discounting measures did not change the pattern of results (see additional data and figures shared on OSF at https://osf.io/htq56/).

## Discussion

Here, we examined whether time, probability, and effort discounting of monetary rewards were related to individual differences in DA function in humans. We found that preferences for shorter time delays, higher probability, and lower physical effort were generally uncorrelated with DA D2-like receptor availability across brain regions in healthy adults.

A meta-analysis comparing correlations between discounting and striatal dopamine function failed to detect a correlation greater than zero. Consistent with Study 1, DA and discounting in healthy groups were unrelated. However, there was heterogeneity dependent on psychopathology, with addiction showing a strong negative relationship to DA. Taken together, these findings suggest that individual differences in D2-like receptors are not reliably associated with discounting in healthy adults. Despite numerous past findings suggesting a role for DA in reward discounting behavior, the present findings raise questions about the specific role of D2-like receptors in discounting.

The difference in correlations between healthy adults and clinical groups in the meta-analysis suggests that individual differences may depend on alterations in striatal DA function. In addictions, striatal D2-like receptor expression is diminished (Volkow et al., 2009), however see (Potenza, 2013) for discussion of mixed findings in pathological gambling potentially due to specific facets of the disorder. This lowered striatal D2-like receptor expression may not be compensated by other features of the DA system such as synthesis capacity, release, re-uptake, or metabolism, which also become dysregulated in addictions (Volkow et al., 2009). As a result, it is possible that effects on temporal discounting emerge when the system is dysregulated.

Dysregulation in different features of the DA system may contribute to non-linear individual differences. The inverted-U hypothesis, for example, has been invoked to characterize individual difference associations between DA and cognition (Vijayraghavan et al., 2007; Cools and D’Esposito, 2011). In this case, changes in D2-like receptors may shift the relative balance in extracellular DA binding with D1-like receptors. Studies have proposed similar inverted-U associations between striatal DA and trait-level sensation-seeking (Gjedde et al., 2010) or fMRI reward signals (Dreher et al., 2008), and cortical DA and delay discounting (Smith and Boettiger, 2012; Elton et al., 2017). The present meta-analytic results revealed little to no association in the healthy range and positive and negative correlations in psychopathology associated with disrupted DA function. An inverted-U relationship driven by dysregulation of striatal DA may account for the differential associations between discounting and D2-like receptors between healthy and clinical groups. Future studies of individuals with a broad range of disruptions in DA are needed to properly test this hypothesis.

Importantly, the measures of baseline D2-like receptor availability were static and cannot describe temporal changes in dopamine signaling related to reward cues. Potentially, individual differences only emerge as a result of temporal dynamics of DA midbrain spiking or DA release (which may also be affected by psychopathology). For example, phasic changes in rodents’ striatal dopamine release vary with discounting behavior (Moschak and Carelli, 2017) and subjective value (Schelp et al., 2017). Phasic changes might better explain individual differences in human reward discounting. For example, value-related fMRI activation linked to the decision process may better capture individual difference associations with baseline DA.

The striking difference in meta-analytic correlation effects between healthy controls and individuals with psychopathology suggests that individual difference findings in clinical samples cannot be reliably generalized to healthy controls, and vice-versa. Disruption of brain function as a result of addiction, ADHD, obesity, and Parkinson’s disease is not limited to a striatal DA abnormality and is more widespread across systems. Alterations in the DA system may interact with changes to broader neural systems. For example, one model of addiction suggests multiple cognitive and motivational corticostriatal circuits interact and compensate for disruptions in glutamatergic and GABAergic prefrontal signaling (Volkow et al., 2011). Disruptions to these circuits may affect the relationship between DA and discounting in addiction (MacKillop et al., 2011). In the context of reward processing, DA release in the striatum impacts cholinergic (Wang et al., 2006), glutamatergic, and GABAergic signaling (Alexander and Crutcher, 1990; Karreman and Moghaddam, 1996). Changes in these other systems may moderate effects of D2-like receptors on discounting, although future studies with direct measures of these system interactions are needed to evaluate this possibility.

Two of the samples in our empirical analysis included age ranges wider than most PET studies of DA. Although age was negatively correlated with D2-like receptor availability, we did not observe age-related associations with discounting in any task. Although prior studies described age differences in discounting (Green et al., 1999; Simon et al., 2010), the lack of an association in the present study is consistent with a recent study of over 23,000 adults which did not identify a correlation between age and time discounting (Sanchez-Roige et al., 2018). Well-documented age-related D2 receptor loss with no changes in discounting behavior is complementary evidence that individual differences in discounting are not likely to be D2-mediated in healthy adults. Controlling for main effects of age did not substantially change any of the results of the primary analyses, suggesting that overall the broad age range of our samples did not account for the lack of effects. However, exploratory analysis of age by D2-like receptor availability interactions revealed that associations between D2-like receptor availability and effort discounting varied across adulthood such that associations were more positive in younger adulthood (particularly in the ventral striatum) and more negative in older adulthood (particularly in the midbrain, where the signal primarily reflects autoreceptors). If replicated, this pattern might suggest that changes in the mesolimbic dopamine system with age have differential impact on effort-based decision making. However, it should be noted that these analyses are based on sample 2 (the only sample that included the effort task), so the within-group analyses of effects are based on relatively few participants. Future research with larger samples across adulthood are needed to better assess the reliability of these effects.

There are several weaknesses of the present studies. Since the finger-pressing requirement for the effort task was not very difficult for participants, additional studies that elicit broader individual differences in preferences are needed to better evaluate associations between D2 receptors and effort discounting.

In the empirical study, we included data from two radiotracers, that have different kinetic properties. Because of this, tracer is confounded with other sample differences. However, in many regions only one tracer contributed data. Furthermore, in primary analyses we included sample as a covariate and observed no significant interactions between sample and D2-like receptor availability in predicting discounting.

The meta-analysis included data from multiple studies using tracers with complementary coverage, but our empirical study was limited to D2-like receptors. Future studies may benefit from comparing multiple measures in the same individuals, for example, D2-like receptors and DAT, the latter of which have been more consistently associated with altering discounting behavior (Wade et al., 2000; van Gaalen et al., 2006; Koffarnus et al., 2011). Although the meta-analysis included one DAT, two DA synthesis, and two DA release effects, radiotracer target did not impact the overall effects in the present analyses, and importantly, analyses restricted to D2 receptors did not impact results. However, given the limited number of effects for most dopamine targets, it is difficult to systematically evaluate potential variation across the dopamine system.

Although the meta-analysis included studies with subject samples varying broadly in clinical status, there was often only one effect per diagnostic group. Importantly, effects from the other psychopathology group that included ADHD, obesity, and PD should be interpreted with caution. Although these groups are similar in that they are impacted by alterations in DA function, there are differences in how DA is dysregulated in each of them (e.g., presynaptic synthesis capacity, DA reuptake, post-synaptic receptor expression) (Madras et al., 2005; Benton and Young, 2016; Kaasinen and Vahlberg, 2017). Grouping of addictions might present issues with respect to illness duration since alterations to DA can exhibit different immediate and long-term changes with drug use (Volkow et al., 2009).

Our meta-analytic results were restricted to temporal discounting, but they were not impacted by the inclusion of correlations for probability and effort discounting tasks.

Unfortunately, there were too few of these different task associations to properly evaluate potential differential effects. Further, the absence of a strong relationship between time, probability, and effort discounting in the empirical data complicates our ability to generalize preferences across tasks. It is possible that the meta-analytic effects observed for time discounting may be different if a greater number of effects for probability and effort were observed. For example, gamblers discount over time but exhibit risk insensitive preferences (Holt et al., 2003), suggesting that probability and time discounting may be different in addiction. Thus, to better characterize specific diagnostic groups affected by alterations in DA function, more studies are needed to evaluate associations with various forms of discounting.

The present findings indicated that individual differences in D2-like receptor availability are not consistently correlated with trait-level individual differences in reward discounting. Our combination of a relatively large empirical study with a meta-analysis adds confidence to the findings and avoids the common weakness of human PET studies, especially individual difference studies, that typically lack statistical power. Future studies specifying the relationship between baseline DA function, temporal dynamics of DA release, and discounting will likely provide additional insight into how dopaminergic control of signaling influences decision preferences in healthy individuals.

## Acknowledgements

This research was supported by National Institute on Aging Pathway to Independence Award R00-AG042596, National Institute on Aging grant R01-AG044838, and National Institute on Drug Abuse grant R21-DA033611. We thank Kevin S. LaBar for comments on portions of the manuscript.

